# Characterizing the Effects of Chronic Cannabis Vapour Exposure and Withdrawal on Cannabinoid Triad, Somatic Signs and Behavioural Network Reorganization in Adult Male Rats

**DOI:** 10.64898/2026.06.12.731896

**Authors:** Abdalla M. Albeely, Hakan Kayir, Richard Quansah-Amissah, Bahar Zali, Selim Karahan, Jaiden Smith, Asim A. Ibrahim, Ahmad Hassan, Sara Hussein, Jude A. Frie, Jibran Y. Khokhar

## Abstract

**Rationale:** Cannabis withdrawal contributes to relapse in individuals with cannabis use disorder, yet preclinical studies have largely focused on withdrawal induced by injected cannabinoids rather than inhaled cannabis, which remains the most common route in humans. The behavioural effects of chronic exposure to vapourized cannabis flower and resulting withdrawal after cessation of exposure remain poorly characterized.

**Objectives:** To determine the behavioural effects of chronic vapourized high-THC cannabis flower exposure on cannabinoid tetrad, somatic withdrawal and behavioural transition networks in rats following both chronic vapour exposure and administration of the cannabinoid receptor 1 (CB1) receptor antagonist SR141716A (rimonabant).

**Methods:** Two studies were conducted using adult male Sprague Dawley rats. The first study (N = 16) exposed rats to either air or vapourized high-THC cannabis flower three times a day for seven days using a Volcano vapourizer, followed by intraperitoneal administration of the CB1 antagonist SR141716A (3 mg/kg). The second study (N = 24) included two air controls and two cannabis groups, with one of each receiving either saline or SR141716A. Behavioural assessments included triad measurements to confirm the cannabis effect, along with withdrawal assessment via a sucrose preference test and somatic signs 30 minutes following rimonabant administration.

**Results:** Repeated cannabis vapour exposure produced reduced locomotor activity, hypothermia, and increased tail-flick latency. Rimonabant administration precipitated withdrawal characterized by increased total withdrawal scores and somatic signs, including blinking, body shakes/tremors, and grooming-related behaviours. Behavioural network analyses revealed substantial reorganization of behavioural transition structure during both chronic cannabis exposure and withdrawal. Chronic cannabis exposure was associated with reduced network modularity, a condensed behavioural repertoire, and altered behavioural centrality measures. At the same time, precipitated withdrawal further increased the influence of exploratory behaviours, particularly sniffing, and reduced the network prominence of locomotor-associated behaviours, such as walking, beyond that detected using conventional behavioural measures alone.

**Conclusion:** Chronic exposure to vapourized cannabis flower followed by CB1 receptor antagonism produces reliable withdrawal symptoms in rats. Behavioural network analyses further reveal that cannabis exposure and withdrawal are both associated with widespread reorganization of behavioural dynamics, suggesting that withdrawal alters not only individual behaviours but also the structure of behavioural transitions. These findings establish a translational model of cannabis withdrawal using inhaled cannabis flower vapour and identify behavioural network analysis as a sensitive approach for characterizing withdrawal-related behavioural states.

## Introduction

Cannabis is one of the most used psychoactive substances in the world and contains over 100 different chemical components, with Δ^9^-tetrahydrocannabinol (THC) being the main psychoactive component. The cannabinoid receptors 1 (CB1) and 2 (CB2) receptors mediate the psychoactive effect of THC as well as other endogenous cannabinoid ligands (Huestis et al., 2001; Pertwee, 2006), and their activation plays a pivotal role in the recreational use of cannabis (Haney, 2022; Huestis et al., 2001).

The high prevalence of cannabis use, coupled with the development of cannabis use disorders (CUD) in 30% of cannabis users and a lack of approved pharmacotherapies for this indication, makes this an important area of study. Withdrawal from cannabis is observed in roughly half of individuals with regular cannabis use or CUD when they suddenly stop using or substantially decrease their consumption of cannabis products containing THC (Connor et al., 2022). While early reports downplayed the presence of cannabis withdrawal, it has been well-established and characterized over the past two decades (Bonnet and Preuss, 2017; Harte-Hargrove and Dow-Edwards, 2012). Nevertheless, withdrawal can be described by a range of symptoms. For instance, in humans, these symptoms, which can occur as early as 24-48 hours after cannabis cessation (Budney et al., 2004), are represented by different somatic signs such as irritability, insomnia, decreased appetite, mood swings and physical discomforts such as shakiness/tremors, sweating, chills, and headache (Connor et al., 2022). Daily users of cannabis also exhibit increased irritability, anxiety, stomach pain and decreased sleep as well as social interaction during cessation (Haney et al., 2003, 2001, 1999). Importantly, when comparing smoked cannabis to oral THC, withdrawal was only observed in individuals smoking cannabis (Hart et al., 2002).

In animals, previous studies have tried to model withdrawal using repeated injections of THC or other cannabinoid agonists and have reported different withdrawal symptoms, including anxiety, irritability, or somatic signs such as tremors, wet-dog shakes and head shakes (Aceto et al., 1995; Beardsley et al., 1986; Brewer et al., 2024; Wilkerson et al., 2019a). Many studies have precipitated cannabinoid withdrawal in animals employing SR141716A (rimonabant), a selective CB1 receptor antagonist, which induces immediate and quantifiable withdrawal symptoms (Cook et al., 1998; Lichtman et al., 2001; Tsou et al., 1995). Administration of rimonabant following prolonged exposure to THC induced distinct somatic withdrawal symptoms, including facial rubs, paw shakes, wet-dog shakes, head shakes and ptosis (Cook et al., 1998; Lichtman et al., 2001). Additionally, Tsou and colleagues reported that rimonabant administration in THC-tolerant rats produced not only somatic withdrawal signs, but also a profound disruption in behavioural organization characterized by rapidly alternating and fragmented sequences of motor behaviours (Tsou et al., 1995). Taken together, these findings demonstrate that cannabinoid withdrawal affects not only the expression of individual withdrawal signs. but may also alter the organization and sequencing of behaviour.

Despite these observations, little work has focused on withdrawal following more ecologically valid inhaled cannabis exposure methods, and behavioural organization during cannabis withdrawal has subsequently received limited quantitative investigation. Considering the pharmacokinetic differences between systemic injections and inhalation, the present study aimed to: 1) characterize withdrawal following chronic exposure to vapourized cannabis flower; and 2) determine whether withdrawal is associated with reorganization of behavioural transition networks in addition to conventional somatic withdrawal signs.

## Methods

### Animals

Animals were pair-housed in polypropylene cages, with free access to chow food and were maintained in a 12-h light cycle. All procedures followed the guidelines described in the Guide to the Care and Use of Experimental Animals (Canadian Council on Animal Care, 1993) and were approved by the Animal Care Committee at the University of Guelph.

### Study 1

Sixteen adult male Sprague Dawley rats (Charles Rivers Laboratories) weighing approximately 350-500 g were randomly assigned to two experimental groups (N = 8/group). The control group (Air+Rim) received hot air (volcano turned on without cannabis flower) while the THC group (Can+Rim) received high-THC cannabis vapour (Truro Wedding Mint, 33.0% THC; Ontario Cannabis Store; 1.5 g) three times a day every 8 hours for 7 days. On day 7, after the last cannabis exposure, rats were evaluated on the triad test (body temperature, locomotor activity, and tail-flick latency) (Zagzoog et al., 2020). On day 8, 12 hours after the final cannabis exposure, both groups received Rimonabant intraperitoneally (i.p.) (3 mg/kg), and 30 minutes later, animals were recorded for 10 minutes to assess somatic signs of withdrawal (gasps, genital licks, foot licks, paw tremors, eye blinks, head and body shakes, chews, ptosis, grooming, and scratches). Animals were also evaluated on the triad test before finishing with a sucrose preference test.

### Study 2

To assess the impact of Rimonabant exposure alone, and to assess whether spontaneous withdrawal could be observed after our vapourized exposure paradigm, we also evaluated a larger second cohort with additional groups. Twenty-four adult male Sprague Dawley rats (Charles Rivers), weighing 350-500 g, were randomly assigned to four experimental groups: Air+Sal (N = 6), Air+Rim (N = 6), Can+Sal (N = 6), and Can+Rim (N = 6). The two control groups were exposed to hot air (as above), and on the last day, one group received i.p.. saline, while the other group received rimonabant (3 mg/kg). In contrast, the THC groups were exposed to high-THC cannabis vapour (Truro Wedding Mint, 33.0% THC; 1.5 g) as above. One group received i.p. saline on the last day, while the other group received rimonabant (3 mg/kg). The same assessments as described in Study 1 were conducted.

### Drugs

Cannabis flower Truro Wedding mint, 33.0% THC, was purchased from the Ontario Cannabis Store (Lot # WM2183). Rimonabant (SR141716A) (RTI International, CAS # 158681-13-1) was solubilized in an 18:1:1 mixture of Saline/ethanol/cremophor and was administered i.p. (3 mg/kg).

### Cannabinoid Triad Test

Rats were tested on the cannabinoid behavioural triad of hyperlocomotion, hypothermia, and antinociception as described (Metna-Laurent et al., 2017) to evaluate the vapourized cannabis-mediated triad effects, and the impact of rimonabant on these effects. Catalepsy was not measured because it has previously not been observed with vapourized cannabis exposure (Marshell et al., 2014; Nguyen et al., 2016).

#### Locomotion

Locomotor activity was assessed in a black acrylic arena (45 × 45 × 45 cm). Rats were habituated to the arena for two days before testing began. Locomotor activity was evaluated at baseline, after the last cannabis exposure and 30 minutes post-rimonabant injection. Rats were evaluated for 10 minutes, and their activity was automatically assessed using Noldus EthoVision XT16 software (Noldus, Leesburg, VA).

#### Hypothermia

Body temperature was measured using a digital microprobe thermometer with a lubricated rectal probe (Model BAT-4, Physitemp Instruments Inc., Clifton, NJ) at baseline, 10 minutes after the last cannabis exposure and 30 minutes post-rimonabant injection.

#### Antinociception

The tail-flick latency assay was used to assess thermal pain sensitivity. Rats were wrapped in a towel, and the distal tip of the rat’s tail (1.5cm) was placed on a tail-flick apparatus heat lamp (Columbus Instruments, Columbus, OH) set to 52 °C. The duration for the rat to display a nociceptive response (tail-flick) was recorded, with a maximum time limit of 30 seconds.

### Evaluation of Somatic Signs of Precipitated Withdrawal

Somatic signs were evaluated by placing each rat into clear acrylic cylindrical chambers (12 x 12 inches) (Frie et al., 2024). The back end of the clear cylinder had a mirror placed facing the camera, allowing for observing the rat when it faces away from the camera. Initially, animals were habituated to the cylinders two days before the start of cannabis exposure for 10 minutes each day. On the test day, 30 minutes after Rimonabant administration, rats were placed into the cylinder and recorded for 10 minutes using a GoPro camera. Videos were re-named and scored by trained observers who were blinded to the experiment setup. Activity in the chamber was assessed for somatic signs of withdrawal, which included blinking, body shakes and tremors, scratching, foot licking, and grooming. Rearing frequency was also assessed using the Noldus EthoVision XT16 software.

### Sucrose Preference Test

The sucrose preference test was used to measure anhedonia (Serchov et al., 2016; Willner et al., 1987). For this test, rats were habituated to 30 mL sipper tubes for two days. Following that, animals were placed on water restriction for two hours before being individually housed and given free access to two 30 mL sipper tubes containing either water or 1.5% sucrose solution for 2h. Bottle positions were counterbalanced between animals to avoid any positional bias. The amount of water and sucrose consumed by each animal was recorded at the end of the second hour.

### Statistical Analysis

Data were analyzed using IBM SPSS Statistics 29 (Armonk, NY, USA) and visualized using GraphPad Prism (San Diego, CA, USA). All Behavioural assessment analyses for Study 1 were performed using Student’s *t*-test. Data from Study 2 were analyzed using a one-way ANOVA across the four treatment groups (Can+Sal, Can+Rim, Air+Sal, and Air+Rim) followed by post hoc comparisons where appropriate. Normality for all data and equality of variance were assessed before statistical analysis using the Shapiro–Wilk test for normality and Levene’s test for equality of variance. The results indicated that the data met the normality and variance homogeneity assumptions, ensuring subsequent statistical analyses.

Network analyses were performed in Python 3.13.9 using NetworkX, SciPy and stats models packages (Hagberg et al., 2008). Initially, eight different behaviours, grooming, jumping, rotation, sniffing, twitching, walking, supported and unsupported rearing, were detected using EthoVision XT v.16.0.1538 (Noldus, Wageningen, NL) Rat Behaviour Recognition Module. Raw data was exported, and the number of transitions between these behaviours was calculated using a custom Python script. Transition counts were expressed as percentages of the total transitions and used as edge weights of network construction. For each animal, a separate directed weighted network was created for each condition (treatment group and experiment time point), with edges retained only when the transition rate exceeded 0.025. For each network, the following global metrics were calculated: density, edge number, total strength, average edge weight, clustering coefficient, transitivity, modularity, number of modules, average path length, reciprocity, and assortativity. For each behavioural node within each network, the strength, clustering, and participation, as well as the following centrality values, were calculated for each animal: degree, betweenness, closeness, Eigenvector, PageRank, and Katz. The difference networks were constructed using the difference values of edge weights. These raw values were used for the statistical evaluations of node-level metrics, but to be able to show them on a single heatmap, a z-score standardization was applied to all groups for each metric for each time point. Rats that did not perform a given behaviour at a given time point were excluded from pooling and averaging for that behaviour, rather than being imputed. Group differences were assessed using two-way repeated measures ANOVA followed by Student’s t-test for post-hoc comparisons. The alpha value was set at the 0.05 level. When only two groups were compared, paired or independent samples t-tests were used as appropriate. Given the large number of simultaneous comparisons across behaviours and node-level metrics, all p-values were corrected for multiple comparisons using the Benjamini-Hochberg false discovery rate (FDR) procedure, with a corrected threshold of q < 0.05 considered statistically significant.

## Results

### Study 1

The studies followed the timeline described in Fig. 1A. Initially, the study evaluated the effect of cannabis flower vapour exposure on cannabis triad measurement at baseline, after the last cannabis exposure, and after rimonabant. No group differences were observed at baseline (Fig. 1B, C, D). After the last vapourized cannabis exposure, the Can group exhibited reduced locomotor activity (t(12) = 3.25, p = 0.007) (Fig. 1B), hypothermia (t(14) = 5.379, p < 0.001) (Fig. 1C), and increased tail-flick latency (t(13) = –3.392, p = 0.005). After Rimonabant, this group only showed increased tail-flick latency (t(13) = –2.728, p = 0.017) (Fig. 1D).

**Figure 1:**
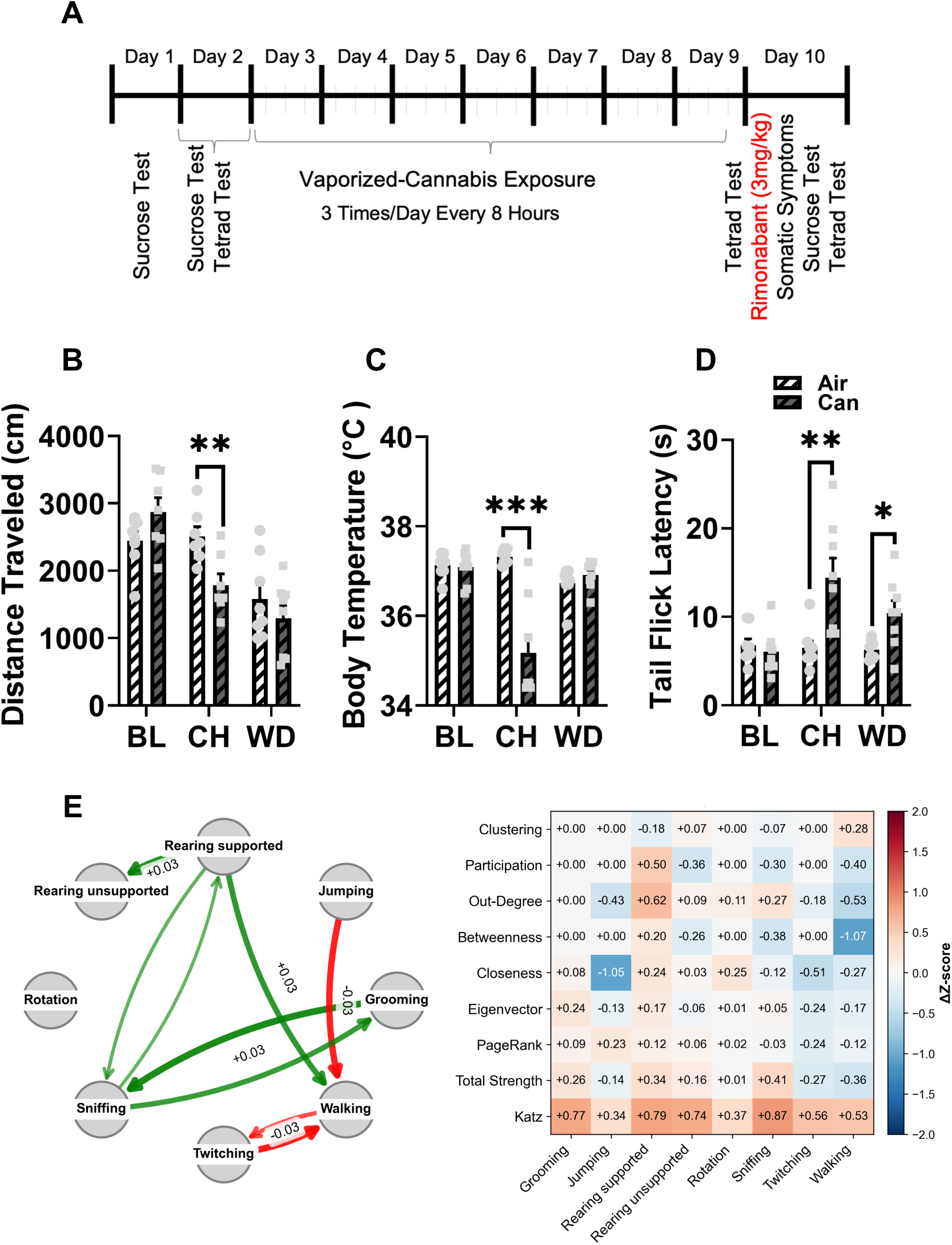
Chronic cannabis exposure induces behavioural and network-level alterations during withdrawal. (A) Experimental timeline showing repeated vapourized cannabis exposure followed by rimonabant-precipitated withdrawal testing. (B–D) Effects of chronic cannabis exposure on locomotor activity (distance travelled), body temperature, and tail-flick latency across baseline (BL), chronic exposure (CH), and withdrawal (WD) timepoints. Chronic cannabis exposure reduced locomotor activity (B) and body temperature (C) while increasing tail-flick latency (D) during both chronic exposure and withdrawal. (E) Behavioural transition network and node-level network analyses during withdrawal. Left: Difference network illustrating altered behavioural transitions in cannabis-exposed animals relative to controls, where green edges indicate increased transitions and red edges indicate decreased transitions. Right: Heatmap showing ΔZ-scored node-level network metrics across behaviours. Cannabis withdrawal was associated with increased exploratory and rearing-related network influence alongside reduced locomotor– and twitching-related centrality measures. Data are presented as mean ± SEM. *p < 0.05, **p < 0.01, ***p < 0.001.

Chronic cannabis vapour exposure altered behavioural transition network organization relative to controls (Fig. 1E). Difference network analyses revealed there were an increase in the total strength (p = 0.024) and transitivity (p = 0.011) of the network, indicating an increased transition among behaviours (Fig. 1E, left). Node-level analyses demonstrated selective alterations in behavioural centrality measures during chronic cannabis exposure (Fig. 1E, right). Cannabis-exposed animals exhibited increased total strength associated with supported rearing and sniffing behaviours (p = 0.007 and 0.010, respectively). In contrast, walking betweenness and strength (p = 0.030 and 0.036, respectively), twitching eigenvector, PageRank, and strength (p = 0.001, 0.012, and 0.002, respectively), and jumping degree centrality values (p = 0.016) reduced in response to chronic cannabis exposure (Fig. 1E, right).

To characterize Rimonabant-precipitated withdrawal in vapourized cannabis flower-exposed rats, somatic withdrawal symptoms were assessed. The Can group showed an increased total withdrawal score (t(13) = –2.181, p = 0.048) (Fig. 2A). Specifically, the Can rat group showed an increased blinking score (t(14) = – 3.287, p = 0.007) (Fig. 2B), body shakes and tremor score (t(13) = –2.217, p = 0.045) (Fig. 2C). Conversely, the Can animals showed reduced scratches and foot licks score (t(13) = 2.424, p = 0.031) (Fig. 2D). Although no significant differences were observed in the grooming score (Fig. 2E), the Can group showed higher grooming duration compared to the Air group (t(14) = 2.586, p = 0.021) (Fig. 2F). No differences were observed in the rearing frequency (Fig. 2G) or in sucrose consumption (Fig. 2H).

**Figure 2:**
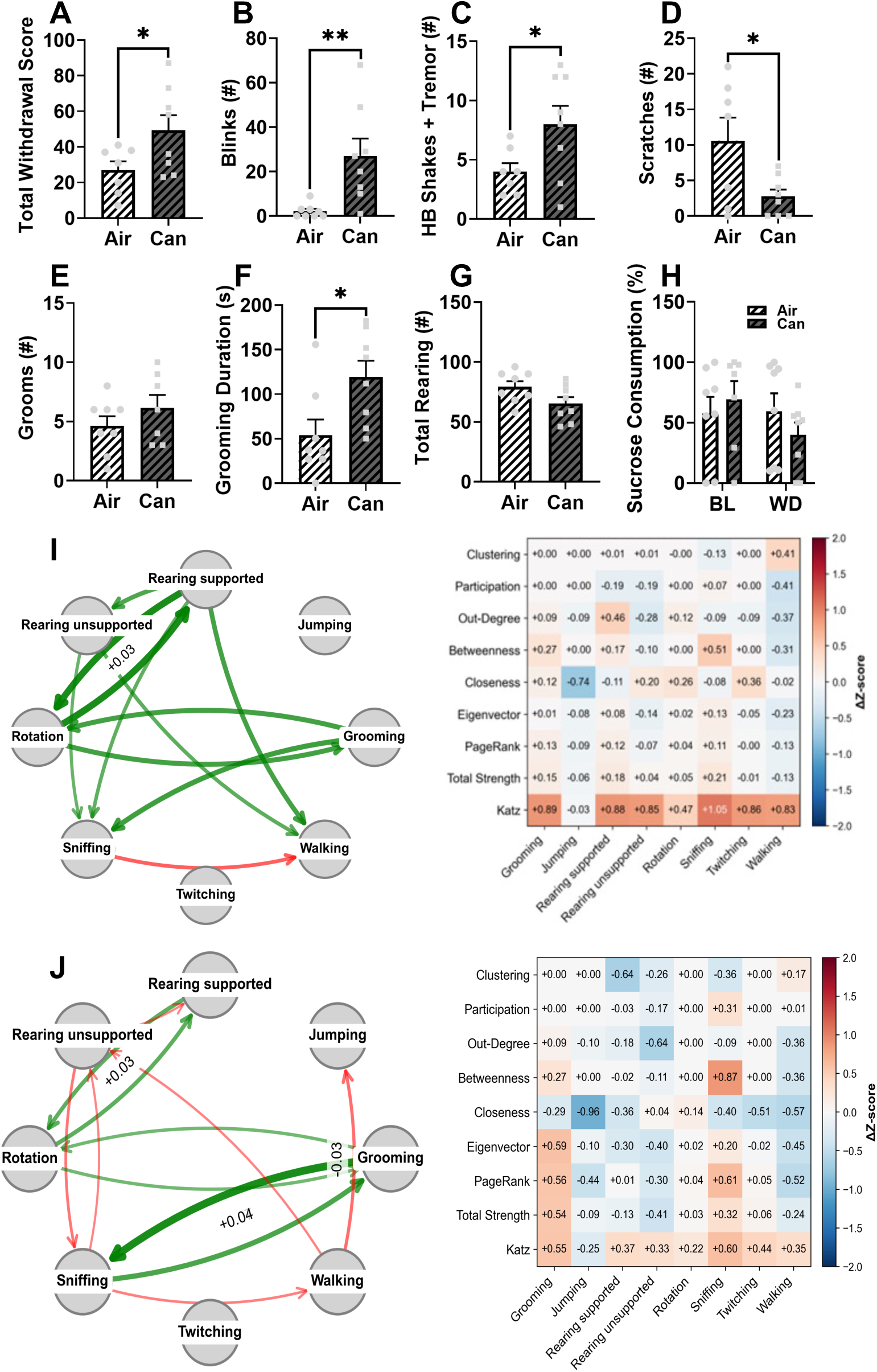
Rimonabant-precipitated withdrawal induces somatic and network-level behavioural reorganization following chronic cannabis exposure. (A–H) Somatic withdrawal signs were measured following rimonabant administration in cannabis-exposed animals. Chronic cannabis exposure increased total withdrawal score (A), blinks (B), head/body shakes with tremor (C), reduced scratches (D) and increased grooming duration (F) compared to air-exposed controls, while no significant differences were observed in grooming frequency (E), rearing frequency (G), or sucrose consumption (H). (I) Behavioural transition network and node-level network analyses during withdrawal relative to controls. Left: Difference network illustrating altered behavioural transitions in cannabis-exposed animals relative to controls, where green edges indicate increased transitions and red edges indicate decreased transitions. Right: Heatmap showing ΔZ-scored node-level network metrics across behaviours. Withdrawal was associated with increased exploratory and rearing-related network influence alongside reduced locomotor-related centrality measures. (J) Direct comparison of behavioural transition organization between chronic cannabis exposure and rimonabant-precipitated withdrawal within cannabis-exposed animals. Left: Difference network illustrating withdrawal-induced alterations in behavioural transitions relative to the chronic exposure state. Right: Heatmap showing ΔZ-scored node-level network metrics during withdrawal relative to chronic cannabis exposure. Withdrawal induced additional restructuring of behavioural transition dynamics and behavioural centrality organization. Data are presented as mean ± SEM. *p < 0.05, **p < 0.01, ***p < 0.001.

During withdrawal, behavioural network organization became further reorganized relative to controls (Fig. 2I). Withdrawal was characterized by a general increase in the strength of the global network (p = 0.031); visually, a widespread increase in transitions among rearing, rotation, grooming, and sniffing-related behaviours, alongside reduced transitions toward walking-related behaviours were observed (Fig. 2I, left). Consistent with these network-level alterations, node-level analyses during withdrawal revealed an increase in sniffing behaviour strength. (Fig. 2I, right). Together, these findings demonstrate widespread reorganization of behavioural transition structure following cannabis exposure and withdrawal. Direct comparison between chronic cannabis exposure and rimonabant-precipitated withdrawal within cannabis-exposed animals did not reveal any significant change at the network level (Fig. 2J). However, node-level analyses demonstrated selective redistribution of behavioural centrality measures during withdrawal, including increased PageRank centrality of grooming (p = 0.026), total strength and degree centrality of unsupported rearing decreased (p = 0.016 and 0.006, respectively). The degree centrality of walking reduced too (p = 0.02). Among remaining behaviours, sniffing was the only behaviour that increased in centrality (Eigenvector centrality, p =0.005), and strength (p = 0.04) (Fig. 2J, right). Together, these findings indicate that precipitated withdrawal induces additional restructuring of behavioural network organization beyond the effects observed during chronic cannabis exposure alone. Full behavioural transition networks and z-scored node-level network matrices for control and cannabis groups are available in Supplementary Fig. S1. Additional behavioural network analyses comparing chronic cannabis exposure and rimonabant-precipitated withdrawal within cannabis-exposed animals are provided in Supplementary Fig. S1-2.

### Study 2

To account for the effects of rimonabant alone, this second study included two groups that received a saline injection on the last day (Air+Sal and Can+Sal). The study followed the timeline shown in Fig. 3A. The triad measurements were evaluated first. When assessing locomotor activity over time, a repeated-measures ANOVA revealed a significant group effect [F(3, 18) = 11.288, p < 0.001]. Following cannabis exposure (before rimonabant injection), Can+Rim animals have displayed reduced locomotor activity compared to Air+Rim (p = 0.007) (Fig. 3B). Can+Sal animals also have showed decreased locomotor activity compared to Air+Sal (p = 0.015) (Fig. 3B). After rimonabant administration, Can+Rim rats exhibited reduced locomotor activity compared to Air+Rim (p = 0.001) (Fig. 3B), while Can+Sal animals showed decreased locomotor activity compared to Air+Sal (p = 0.017) (Fig. 3B).

**Figure 3:**
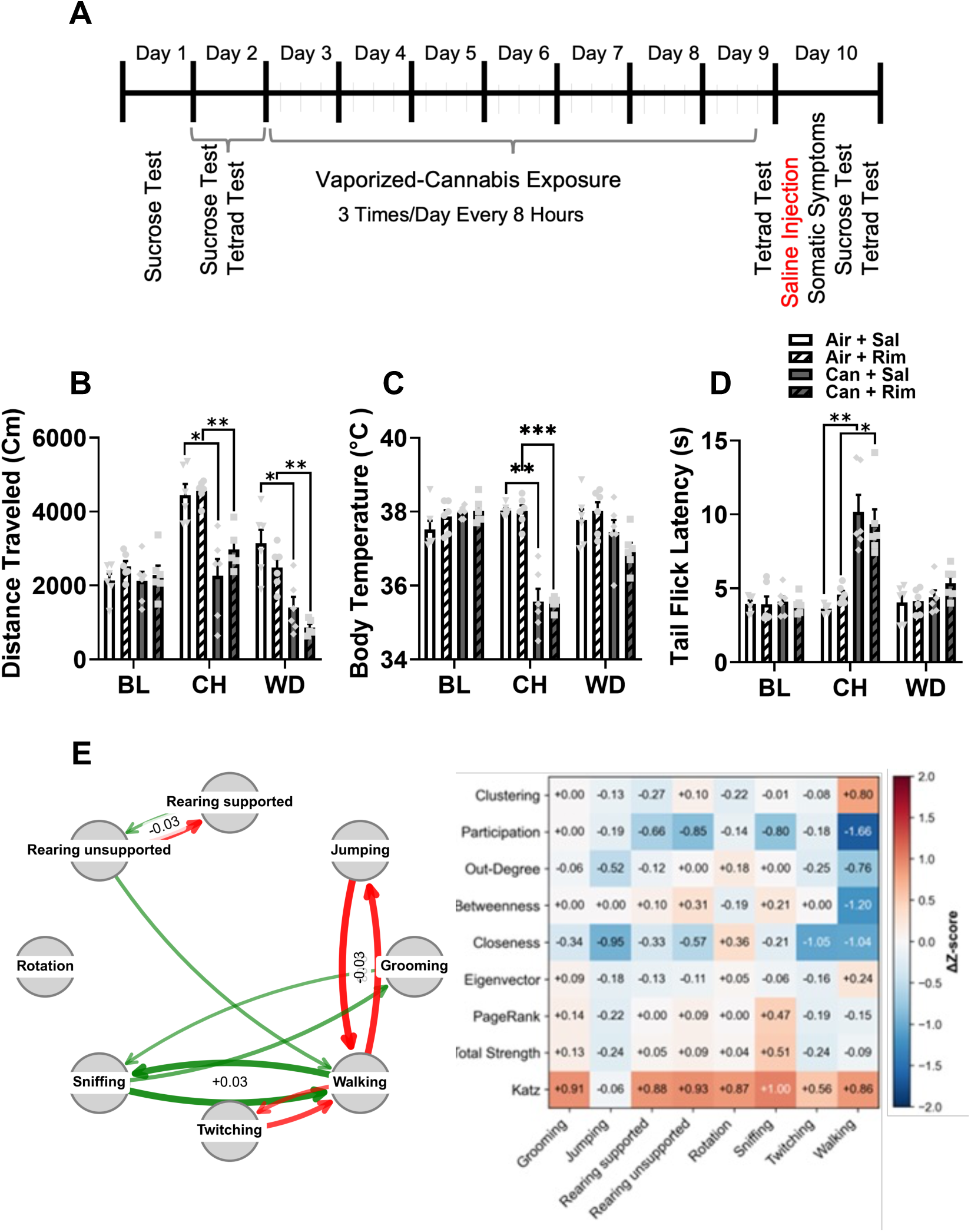
Chronic cannabis exposure and precipitated withdrawal alter behavioural and network organization, with limited effects of rimonabant or cannabis exposure alone on withdrawal. Experimental timeline showing repeated vapourized cannabis exposure followed by saline or rimonabant administration. (B–D) Effects of chronic cannabis exposure and rimonabant (or vehicle) administration on locomotor activity (distance travelled), body temperature, and tail-flick latency across baseline (BL), chronic exposure (CH), and withdrawal (WD) timepoints. Cannabis-exposed animals exhibited reduced locomotor activity (B) and body temperature (C), alongside increased tail-flick latency (D) following chronic exposure. (E) Behavioural transition network and node-level network analyses during chronic cannabis exposure. Left: Difference network illustrating altered behavioural transitions in cannabis-exposed animals relative to air-exposed controls, where green edges indicate increased transitions and red edges indicate decreased transitions. Right: Heatmap showing ΔZ-scored node-level network metrics across behaviours. Chronic cannabis exposure was associated with increased exploratory and grooming-related network influence alongside reduced locomotor– and jumping-related centrality measures. Data are presented as mean ± SEM. *p < 0.05, **p < 0.01, ***p < 0.001.

With body temperature, a significant group effect was observed [F(3,20) = 53.599, p < 0.001] following the last vapourized cannabis exposure. The Can+Sal group showed a reduction in body temperature compared to the Air+Sal (P = 0.003) (Fig. 3C). The Can+Rim also showed lower body temperature compared to the Air+Rim (p <0.001) groups (Fig. 3C). Tail-flick latency showed a significant group effect [F(3,20) = 18.033, p < 0.001; ANOVA] following the last cannabis exposure, with the Can+Sal group showing increased tail-flick latency compared to the Air+Sal (p = 0.009) (Fig. 3D). The Can+Rim group also showed elevated tail-flick latency compared to the Air+Rim (p = 0.016) (Fig. 3D).

Behavioural transition network organization remained altered during chronic cannabis exposure and following rimonabant-precipitated withdrawal (Fig. 3E). During chronic cannabis exposure, difference network analyses revealed the edge number of the network decreased while the average edge weight increased (p = 0.02 and 0.009, respectively), indicating a less diverse network using the remaining behaviour nodes more often. The average path between behaviours increased (p = 0.039), which supports the decreased efficiency of the network. Further supporting the findings that a few dominant behaviours became strongly connected, the modularity of the network decreased (p < 0.001) and the assortativity increased (p = 0.008) (Fig. 3E, left). Node-level analyses demonstrated selective alterations in behavioural network organization during chronic cannabis exposure, mostly related to changes in transitions of walking, jumping, twitching and sniffing behaviours (Fig. 3E, right). The clustering value of walking behaviour increased (q = 0.049) while its participation decreased (q = 0.025), indicating its increased integration with a smaller number of behaviours. Moreover, degree, betweenness, and closeness centralities of walking were reduced (q = 0.043, 0.035, and < 0.016), showing a loosening role of walking as a connector or mediator in the global network (Fig.3E, right). On the other hand, the strength of sniffing increased (q = 0.035) while it decreased for jumping and twitching (q = 0.025 and 0.033, respectively), shifting the sniffing behaviour to a more transitional place in the condensed global network. Participation of sniffing and unsupported rearing decreased (q = 0.014 for both), showing the decrease in their bridging role among diverse behaviours (Fig. 3E, right).

When characterizing the somatic symptoms of withdrawal in this study, an ANOVA for the total withdrawal score showed a significant group effect [F(3,17) = 9.745, p = 0.001]. Both Can+Sal and Can+Rim groups show an increased total withdrawal score compared to the Air+Rim group (p = 0.012 and 0.007, respectively, but not from Air+Sal group) (Fig. 4A). In the grooming score, an ANOVA showed a significant group effect [F(3,19) = 4.689, p = 0.013], with the Can+Rim showing an increased grooming score compared to both the Air+Sal group (p = 0.047), and the Air+Rim group (p = 0.025) (Fig. 4E). When analyzing rearing frequency, a significant group effect was observed [F(3,20) = 3.400, p = 0.38], with the Can+Rim group exhibiting lower rearing frequency than the Can+Sal group (P = 0.018) (Fig. 4G). No other significant differences were observed in shakes, scratches, grooming duration, or sucrose preference (Fig. 4B, C, D, F, and G).

**Figure 4:**
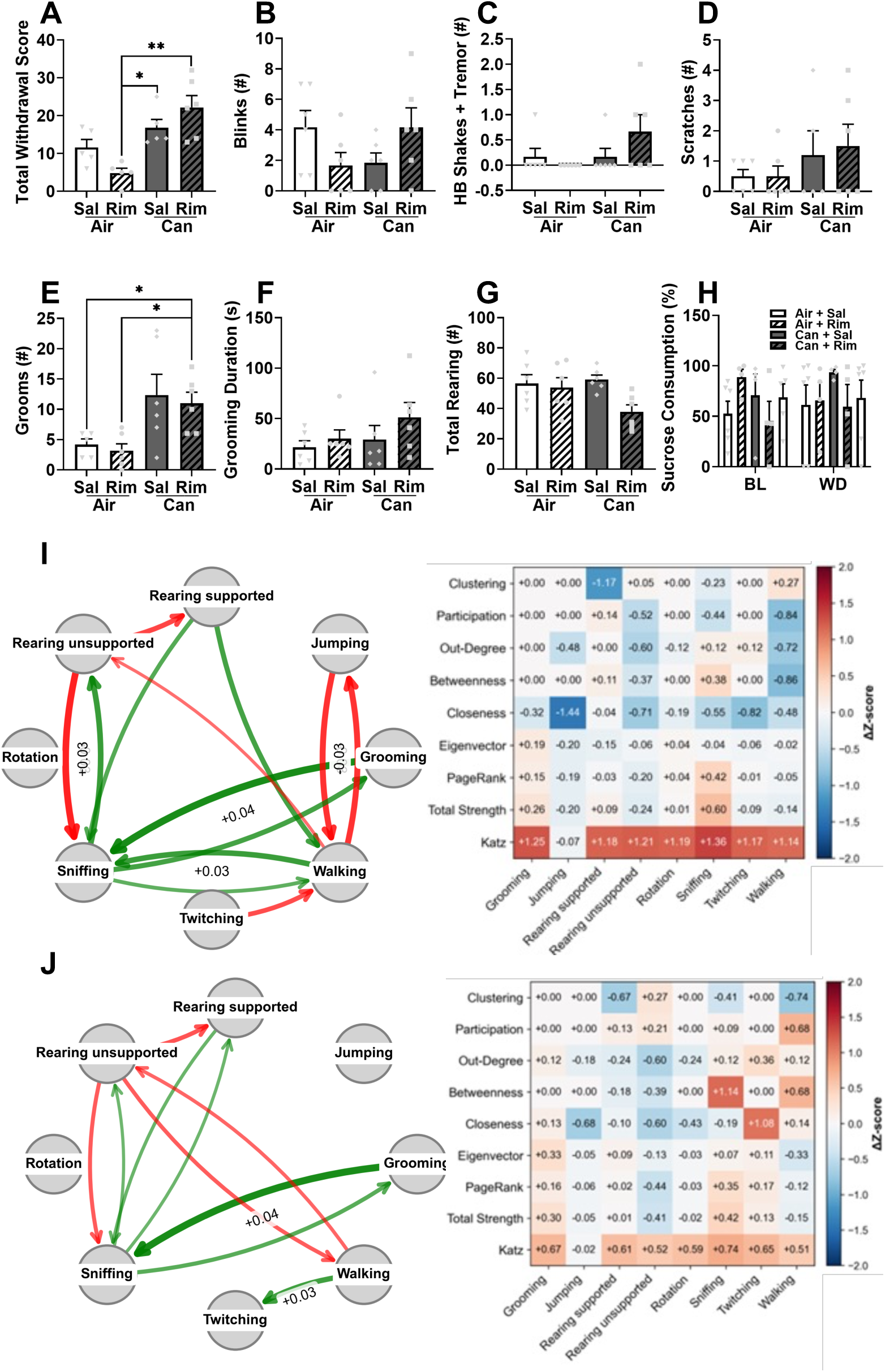
Behavioural and network reorganization during spontaneous and precipitated cannabis withdrawal. (A–H) Somatic withdrawal signs were measured following saline or rimonabant administration in cannabis-exposed and air-exposed animals. Cannabis-exposed animals exhibited increased total withdrawal score and grooming frequency relative to controls, while reduced rearing frequency was observed following rimonabant administration. No significant differences were observed in blinks, head/body shakes with tremor, scratches, grooming duration, or sucrose consumption. (I) Behavioural transition network and node-level network analyses during rimonabant-precipitated withdrawal relative to controls. Left: Difference network illustrating altered behavioural transitions in cannabis-exposed animals relative to controls, where green edges indicate increased transitions and red edges indicate decreased transitions. Right: Heatmap showing ΔZ-scored node-level network metrics across behaviours. Withdrawal was associated with increased exploratory and rearing-related network influence alongside reduced locomotor-related centrality measures. (J) Direct comparison of behavioural transition organization between chronic cannabis exposure and rimonabant-precipitated withdrawal within cannabis-exposed animals. Left: Difference network illustrating withdrawal-induced alterations in behavioural transitions relative to the chronic exposure state. Right: Heatmap showing ΔZ-scored node-level network metrics during withdrawal relative to chronic cannabis exposure. Precipitated withdrawal induced additional restructuring of behavioural transition dynamics and node-level behavioural organization. Data are presented as mean ± SEM. *p < 0.05, **p < 0.01.

Following rimonabant-precipitated withdrawal, weights of the behavioural network showed a significant difference among the groups [F(3,20) = 4.238; p = 0.018] (Fig. 4I). Post hoc analyses indicated a difference between Can+Rim vs Air+Rim (p = 0.015). Further comparisons indicated that modularity and edge number decreased in Can+Rim group (p= 0.011 and 0.018, respectively) compared to Air+Rim group. Thus, the transitions between the behaviours were reduced, and the network used fewer behaviours during rimonabant precipitated withdrawal. Consistent with these transition-level alterations, node-level analyses following rimonabant administration revealed persistent increases in Katz centrality across all the behaviours except jumping (q values < 0.05) (Fig. 4I, right). This global increase in Katz centrality reflects a repetitive shift through indirect pathways, and increased repetition among all these behaviours.

Direct comparison between chronic cannabis exposure and rimonabant-precipitated withdrawal within the same group of animals revealed additional reorganization of behavioural transition networks following withdrawal induction (Fig. 4J). Network level analyses indicated that, relative to the chronic exposure state, withdrawal was associated with a decrease in clustering coefficient (p = 0.036, paired t test), indicating a looser integration of this group of behaviours (Fig. 4J, left). Node-level analyses further demonstrated selective redistribution of behavioural centrality measures during withdrawal, including reduction in degree centrality of unsupported rearing (p = 0.011), and increased degree centrality of twitching (p = 0.025) (Fig. 4J, right). Thus, the role of unsupported rearing behaviour in the integration of other behaviours switched to twitching during precipitated withdrawal compared to the chronic exposure state. Together, these findings indicate that precipitated withdrawal induces additional restructuring of behavioural transition organization beyond the effects observed during chronic cannabis exposure alone. Full behavioural networks and z-scored node-level matrices are provided in Supplementary Fig. S3. Additional induced and spontaneous withdrawal network analyses are shown in Supplementary Fig. S3-5.

## Discussion

The objective of this study was to evaluate whether withdrawal can be precipitated in animals exposed to vapourized cannabis flower. Two experiments, for the first time, revealed that rats displayed rimonabant-precipitated cannabis withdrawal following seven days of exposure to vapourized cannabis three times daily.

To characterize the effects of repeated cannabis vapour exposure, rats underwent a cannabis triad test. In both studies, the observed increases in tail-flick latency, as well as reduction in locomotor activity and body temperature following vapourized cannabis exposure, are consistent with previous reports on the effect of THC, reflecting the well-documented properties of cannabinoids known to modulate motor function, temperature and activity levels (Baron, 2018; Crippa et al., 2010; Monti, 1977). Furthermore, marked reductions in body temperature are consistent with the published findings on vapourized THC exposure (Taffe et al., 2015). Multiple mechanisms may underlie cannabis-induced hypothermia. For instance, serotonin depletion was shown to reduce THC-induced hypothermia, while serotonin reuptake inhibition potentiates the hypothermic response (Davies and Graham, 1980). Additionally, the endocannabinoid system in the hypothalamus, a critical brain region in thermoregulation, may also contribute to reduced body temperature via transient receptor potential vanilloid 1 (TRPV1) (Wenger and Moldrich, 2002). Endocannabinoids like anandamide (AEA) and 2-arachidonoylglycerol (2-AG) are endogenous agonists of the TRPV1 receptor (Ross, 2003). Although THC has minimal efficacy at TRPV1 receptors, other cannabinoids, such as cannabidiol and cannabinol, can activate and desensitize the TRPV1 receptor, leading to hypothermia (DeVuono and Parker, 2020; Rawls and Benamar, 2011). This interaction between TRPV1 receptors and the endocannabinoid system within the hypothalamus may also contribute to the overall hypothermic effect observed in this study. Importantly, administration of the CB1 receptor antagonist rimonabant attenuated cannabis-induced hypothermia in Study 1, supporting the interpretation that this effect is primarily mediated by CB1 receptor activation. CB1 receptors are densely expressed in the hypothalamus, where they modulate thermoregulatory circuits, and blockade of these receptors has been shown to prevent or reverse THC-induced hypothermia in previous studies (Rawls et al., 2002; Wenger and Moldrich, 2002). However, in Study 2, hypothermia persisted in THC-exposed animals regardless of rimonabant treatment, suggesting that additional mechanisms, including sustained cannabinoid signalling or non-CB1 receptor pathways, may contribute to the temperature reductions observed following repeated vapourized cannabis exposure. Together, these findings indicate that cannabis-induced hypothermia in this model is largely CB1-dependent but may also involve secondary modulatory systems.

Analysis of the somatic signs of withdrawal in studies 1 and 2 revealed that the groups exposed to cannabis vapour showed increased total withdrawal scores after withdrawal precipitation by rimonabant. The observed withdrawal symptoms were consistent with previous studies employing repeated THC administration followed by CB1 receptor antagonism (Aceto et al., 1995; Cook et al., 1998; Lichtman et al., 2001; Tsou et al., 1995). Consistent with previous reports, cannabis-exposed animals displayed increased tremors, body shakes, and grooming-related behaviours following rimonabant administration. Tremors have been identified as a reliable indicator of cannabis withdrawal in both rodents (Aceto et al., 1995; Cook et al., 1998; Tsou et al., 1995) and monkeys (Kaymakcalan, 1973), while head shaking and scratching have been reported as common rimonabant-precipitated withdrawal symptoms in mice and rats (Aceto et al., 1995; Lichtman et al., 2001; Tsou et al., 1995). Similarly, the increased grooming score or duration observed in this study is also consistent with previous findings reporting excessive grooming in rats during abrupt, precipitated withdrawal (Pertwee, 1992). Together, these findings indicate that withdrawal induced following chronic exposure to vapourized cannabis flower produces a behavioural profile that closely resembles that reported previously following repeated THC administration.

Withdrawal-related behaviours were most evident following CB1 receptor antagonism, whereas cannabis-exposed animals that received saline displayed relatively fewer detectable withdrawal symptoms. This suggests that seven days of cannabis vapour exposure was sufficient to produce neuroadaptations that could be revealed by antagonist challenge but may be insufficient to generate robust spontaneous withdrawal under the behavioural measures employed here. This finding is consistent with previous work demonstrating that, despite clear evidence of THC-induced locomotor suppression and tolerance development, spontaneous withdrawal following repeated THC exposure in rats is often mild and difficult to detect using conventional behavioural measures (Wilkerson et al., 2019). For example, Wilkerson and colleagues reported no significant changes in voluntary wheel-running behaviour during spontaneous withdrawal following repeated THC administration, suggesting that spontaneous withdrawal symptoms may be subtle relative to those observed during precipitated withdrawal paradigms (Hickey et al., 2026; Wilkerson et al., 2019). Together, these findings support the utility of antagonist-precipitated withdrawal models for revealing dependence-related behavioural alterations that may otherwise remain difficult to detect.

Notably, Tsou and colleagues reported that precipitated withdrawal was characterized not only by somatic signs such as grooming, wet-dog shakes, and tremor-like movements, but also by a “disorganized pattern of constantly changing brief sequences of motor behaviours” (Tsou et al., 1995). Recent advances in automated behavioural analysis pipelines, such as those using pose estimation (DeepLabCut) and active learning frameworks (A-SOiD) in LUPE chambers, have begun to address the challenges of quantifying subtle and rapid withdrawal behaviours in opioid models (Cohen et al., 2025). Here, to further characterize the cannabis withdrawal phenotype and identify alterations in behavioural organization that conventional behavioural measures may not capture, we applied a network analysis approach based on behavioural transition rates (Ashiquzzaman et al., 2025; Hsu and Yttri, 2021). Our network-based approach similarly extends beyond isolated somatic sign counts by quantifying reorganization of behavioural transition dynamics, revealing condensed and less flexible behavioural repertoires during cannabis exposure and withdrawal. These behavioural network analyses revealed that chronic cannabis exposure and withdrawal were associated with substantial reorganization of behavioural transition dynamics. Consistent with this concept, the present findings demonstrate that withdrawal does not simply alter isolated behaviours but instead reorganizes the structure and flow of behavioural dynamics across the network. Our study further extends this concept to the effects of chronic cannabis exposure. Across both studies presented here, exploratory behaviours, including sniffing, exhibited increased transition probability and network influence. In contrast, locomotor-associated behaviours, such as walking, displayed reduced network integration and centrality. Both studies consistently showed a reduced influence of jumping and twitching behaviours after chronic exposure. The second study clearly showed that chronic cannabis exposure condenses the behaviour transition network by creating fewer but stronger connections with reduced modularity, suggesting that cannabis exposure progressively narrows the behavioural repertoire rather than simply reducing individual behaviours. These findings suggest that cannabis withdrawal may shift behavioural organization away from coordinated locomotor behavioural flow toward more fragmented and exploratory behavioural states. Notably, especially in the second study, Katz centrality was consistently increased across withdrawal-related conditions, particularly among exploratory and rearing-associated behaviours. Because Katz centrality reflects both direct and indirect influence within a network, these findings suggest that withdrawal-related behavioural states become more broadly interconnected and propagate more strongly across behavioural sequences. Together, these findings suggest that withdrawal-related behavioural organization becomes increasingly dominated by recurrent exploratory behavioural states, potentially reflecting reduced behavioural flexibility and more constrained behavioural sequencing supported by several network-level findings. These global network changes indicate that the behavioural repertoire became simultaneously narrower, less efficiently connected, and more segregated into fewer dominant transition pathways, collectively reflecting a loss of the flexible modular organization that normally enables smoother transition among behavioural states.

Although several network-level alterations were consistently observed in both studies, the specific behavioural manifestations of withdrawal differed between experiments. In the first study, blinking and head-body shakes and tremors were the dominant withdrawal-related behaviours, whereas grooming-related behaviours were more prominent in the second study. This variability suggests that cannabis withdrawal may not be expressed through a single stereotyped behavioural pattern but rather through multiple withdrawal phenotypes that differ across individuals. Such variation may arise from intrinsic biological differences that influence how animals respond to chronic cannabinoid exposure and withdrawal. Supporting this interpretation, human cannabis users reported substantial heterogeneity in the expression of withdrawal symptoms, including anxiety, irritability, and changes in appetite (Mennes et al., 2009). Together, these findings suggest that variability may be an inherent feature of the cannabis withdrawal phenotype rather than simply experimental noise.

While the current study provides valuable insights into cannabis withdrawal, several limitations should be acknowledged. Firstly, the study assessed withdrawal at only a one-time point following the last cannabis exposure. However, the temporal dynamics of withdrawal symptoms may vary, and more detailed time-course assessments could provide valuable insights into the effects of vapourized cannabis withdrawal. Secondly, the study primarily used one type of cannabis, potentially overlooking the diversity in the constituents of cannabis and their impact on the neurobiological underpinnings of withdrawal symptoms. Given the wide variety of cannabis chemovars and their distinct pharmacological properties, future studies should consider different types to better understand the spectrum of withdrawal symptoms and whether other types might elicit differing levels of withdrawal (Baron, 2018; Pertwee, 2006; Russo, 2011). Furthermore, while the study emphasized different behavioural outcomes, investigating additional measures that may be disrupted during withdrawal, such as sleep patterns, blood pressure, heart rate, and changes in oscillatory activity, could offer valuable insights (Connor et al., 2022; Jenkins et al., 2021; Kolla et al., 2022; Luciani et al., 2020). While telemetry tools can enable monitoring of these physiological parameters, offering real-time data on their fluctuation during withdrawal, electrophysiological measurements, such as local field potentials, can provide critical insight into brain function and connectivity during this process. Integrating telemetry tools to monitor sleep stages, with electrophysiological methods, could provide a more comprehensive understanding of cannabis withdrawal. Moreover, the study employed male rats exclusively, underscoring the need for future research to explore potential sex differences in cannabis withdrawal symptoms. Notably, women have reported more withdrawal symptoms than men, as well as gastrointestinal issues such as nausea and stomach pain (Bonnet and Preuss, 2017; Harte-Hargrove and Dow-Edwards, 2012; Kesner et al., 2022; Marusich et al., 2014). Future studies will incorporate both male and female rats in assessing withdrawal symptoms and the mechanisms underlying these symptoms.

In conclusion, the results of this study provide valuable insights into withdrawal symptoms after exposure to inhaled vapourizd cannabis flower in rats. This helped characterize a platform that can now be used for future mechanistic research as well as testing novel therapeutics. Additionally, the behavioural network analyses used here also prove useful for in-depth assessment of withdrawal from cannabis and other drugs in preclinical models. Further research is needed to explore the neural mechanisms underpinning cannabis withdrawal, with potential implications for developing targeted interventions to alleviate withdrawal symptoms in individuals with cannabis use disorder.

## Data Availability

The datasets generated and analyzed during the current study are available from the corresponding author upon reasonable request.

## Authors Contribution

A.M.A conceived the study, performed the experiments, analyzed the data, and wrote the first draft of the manuscript. HK performed the behavioural network analyses, contributed to data analysis, and assisted with manuscript editing. RQA contributed to the study conception and experimental design.B.Z, S.K, J.S, A.S, A.A.I. and S.H. assisted with behavioural scoring and data collection. J.A.F. contributed to the conceptual development of the study and provided input on experimental design. J.Y.K. conceived and supervised the study, secured funding, contributed to study design and data interpretation, and critically revised the manuscript. All authors reviewed, edited, and approved the final version of the manuscript.

## Supplementary Figure Legends

**Supplementary Fig. S1. behavioural transition networks and node-level network metrics in air and cannabis exposed animals during chronic cannabis exposure and withdrawal from Study 1**. (A–D) Full behavioural transition networks and corresponding node-level heatmaps across chronic exposure and withdrawal timepoints in air– and cannabis-exposed animals. Panels show Chronic – Air group (A), Chronic: Can group (B), Withdrawal: Air group (C), and Withdrawal: Can group (D) conditions. Left: Behavioural transition networks illustrating the strength and directionality of transitions between behavioural states, where edge thickness reflects transition strength. Node size corresponds to behavioural connectivity within the network. Right: Heatmaps showing mean z-scored node-level network metrics across behaviours. Chronic cannabis exposure and withdrawal altered behavioural transition structure and node-level network organization, particularly within exploratory-, grooming-, and locomotor-associated behaviours.

**Supplementary Fig. S2. Behavioural transition networks and node-level network metrics in cannabis-exposed animals during chronic exposure and withdrawal from Study 1**. (A–B) Full behavioural transition networks and corresponding node-level heatmaps in cannabis-exposed animals during the chronic exposure (A) and withdrawal (B) timepoints. Left: Behavioural transition networks illustrating the strength and directionality of transitions between behavioural states, where edge thickness reflects transition strength. Node size corresponds to behavioural connectivity within the network. Right: Heatmaps showing mean z-scored node-level network metrics across behaviours. Withdrawal altered behavioural transition structure and redistributed node-level network organization relative to the chronic exposure state, particularly within exploratory-, grooming-, and locomotor-associated behaviours.

**Supplementary Fig. S3. Behavioural transition networks and node-level network metrics across chronic exposure and rimonabant-precipitated withdrawal conditions from Study 2**. (A–D) Full behavioural transition networks and corresponding node-level heatmaps across chronic exposure and withdrawal conditions in Air+Rim and Can+Rim animals. Panels show Chronic condition for Air+Rim group (A), Chronic condition for Can+Rim group (B), the Withdrawal condition for Air+Rim group(C), and Withdrawal condition for Can+Rim group (D) conditions. Left: Behavioural transition networks illustrating the strength and directionality of transitions between behavioural states, where edge thickness reflects transition strength. Node size corresponds to behavioural connectivity within the network. Right: Heatmaps showing mean z-scored node-level network metrics across behaviours. Chronic cannabis exposure and rimonabant-precipitated withdrawal altered behavioural transition structure and redistributed node-level network organization, particularly within exploratory-, grooming-, and locomotor-associated behaviours.

**Supplementary Fig. S4. Study 2 full behavioural transition networks and node-level network metrics in cannabis-exposed animals during chronic exposure and rimonabant-precipitated withdrawal.** (A–B) Full behavioural transition networks and corresponding node-level heatmaps in Can+Rim animals during the chronic exposure (A) and withdrawal (B) timepoints. Left: Behavioural transition networks illustrating the strength and directionality of transitions between behavioural states, where edge thickness reflects transition strength. Node size corresponds to behavioural connectivity within the network. Right: Heatmaps showing mean z-scored node-level network metrics across behaviours. Rimonabant-precipitated withdrawal altered behavioural transition structure and redistributed node-level network organization relative to the chronic exposure state, particularly within exploratory-, grooming-, and locomotor-associated behaviours.

**Supplementary Fig. S5. Study 2 full behavioural transition networks and node-level network metrics in Can+Sal animals during chronic exposure and spontaneous withdrawal.** (A–B) Full behavioural transition networks and corresponding node-level heatmaps in Can+Sal animals during the chronic exposure (A) and withdrawal (B) timepoints. Left: Behavioural transition networks illustrating the strength and directionality of transitions between behavioural states, where edge thickness reflects transition strength. Node size corresponds to behavioural connectivity within the network. Right: Heatmaps showing mean z-scored node-level network metrics across behaviours. (C) Difference network and ΔZ-scored node-level heatmap comparing spontaneous withdrawal relative to the chronic exposure state in Can+Sal animals. Green edges indicate increased transitions, and red edges indicate decreased transitions during withdrawal. Spontaneous withdrawal altered behavioural transition structure and redistributed node-level network organization, particularly within exploratory-, grooming-, and locomotor-associated behaviours.

