## Supplementary figures and images for "Characterizing the Effects of Chronic Cannabis Vapour Exposure and Withdrawal on Cannabinoid Triad, Somatic Signs and Behavioural Network Reorganization in Adult Male Rats"

### Supplemental Figures

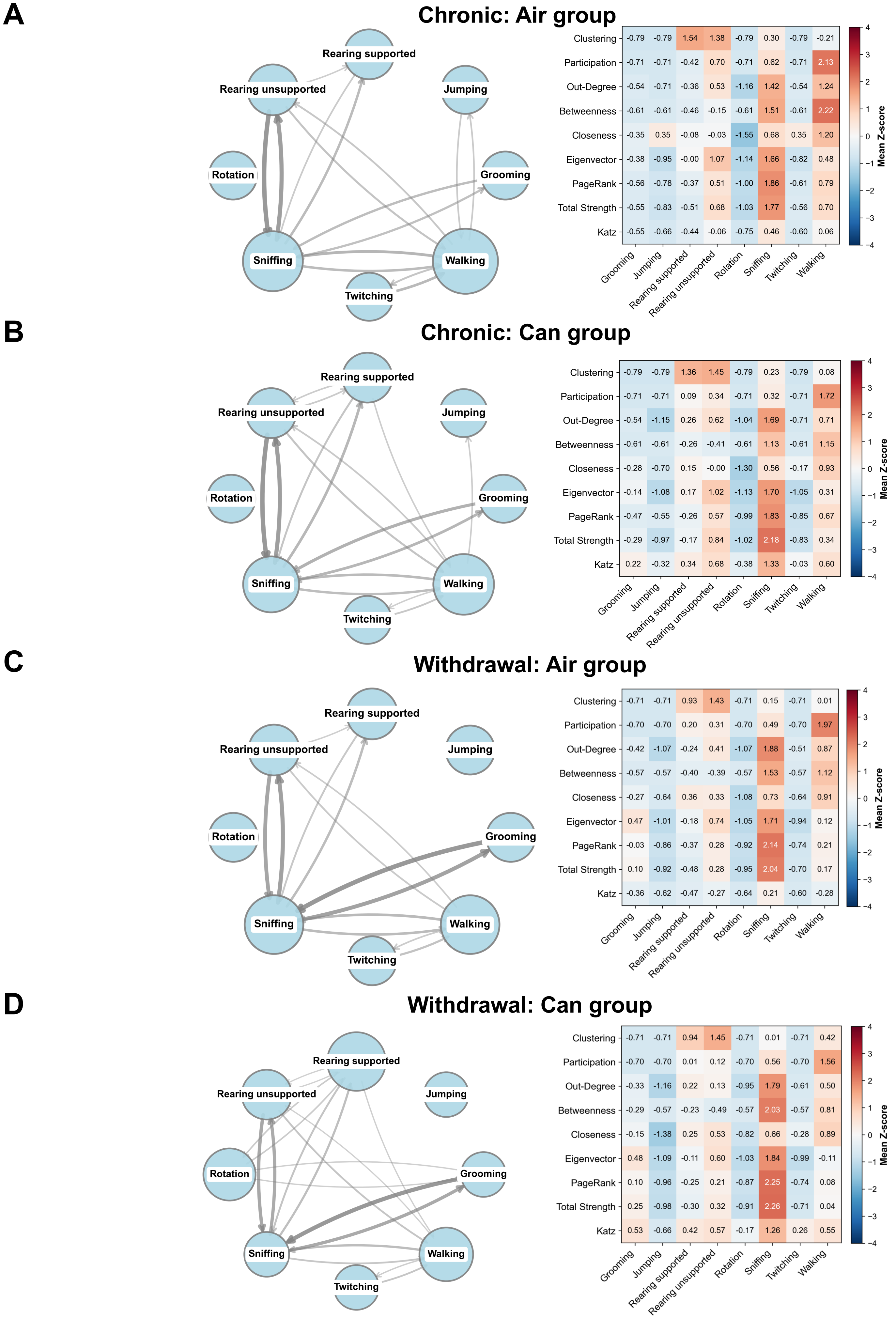

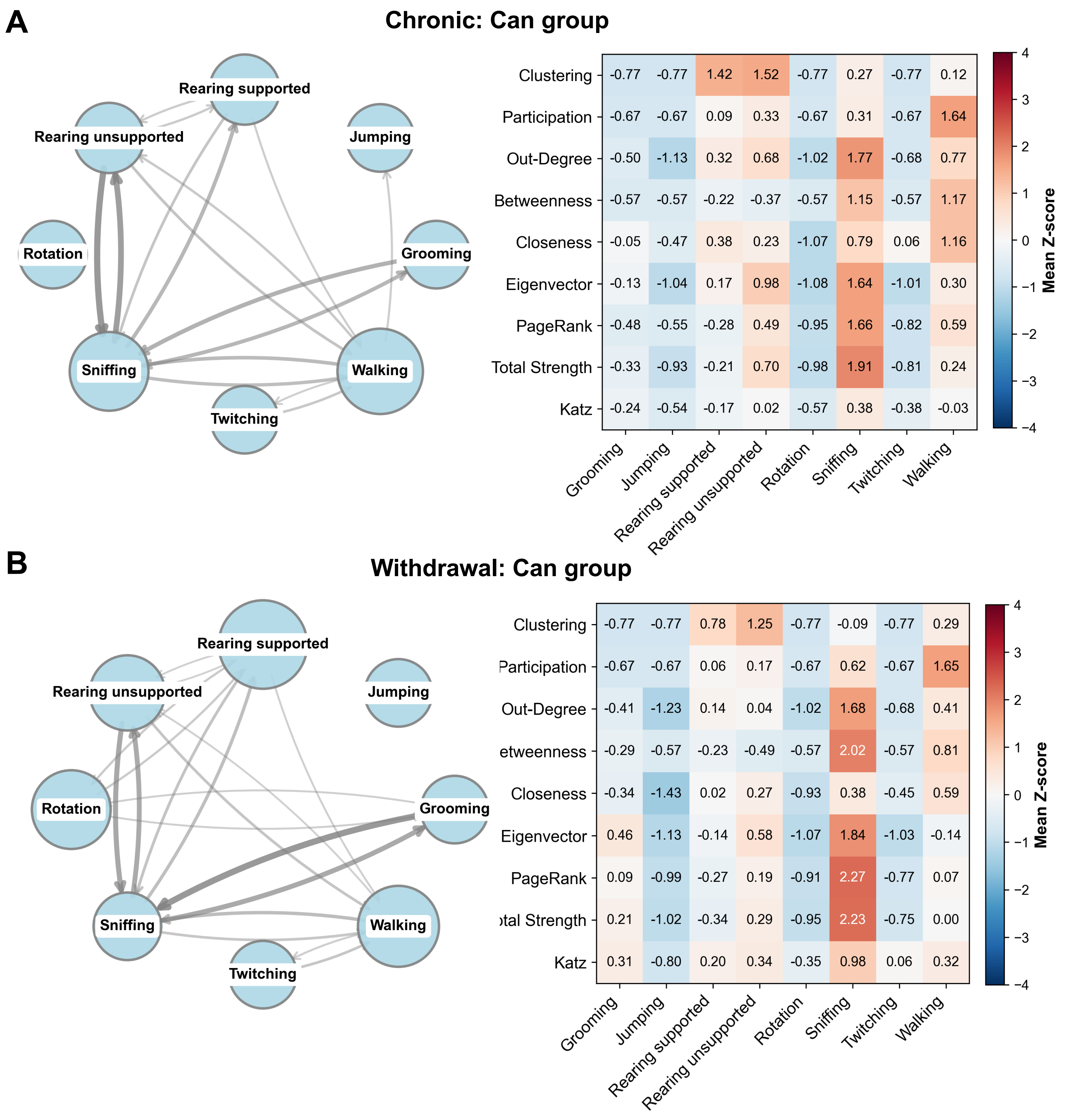

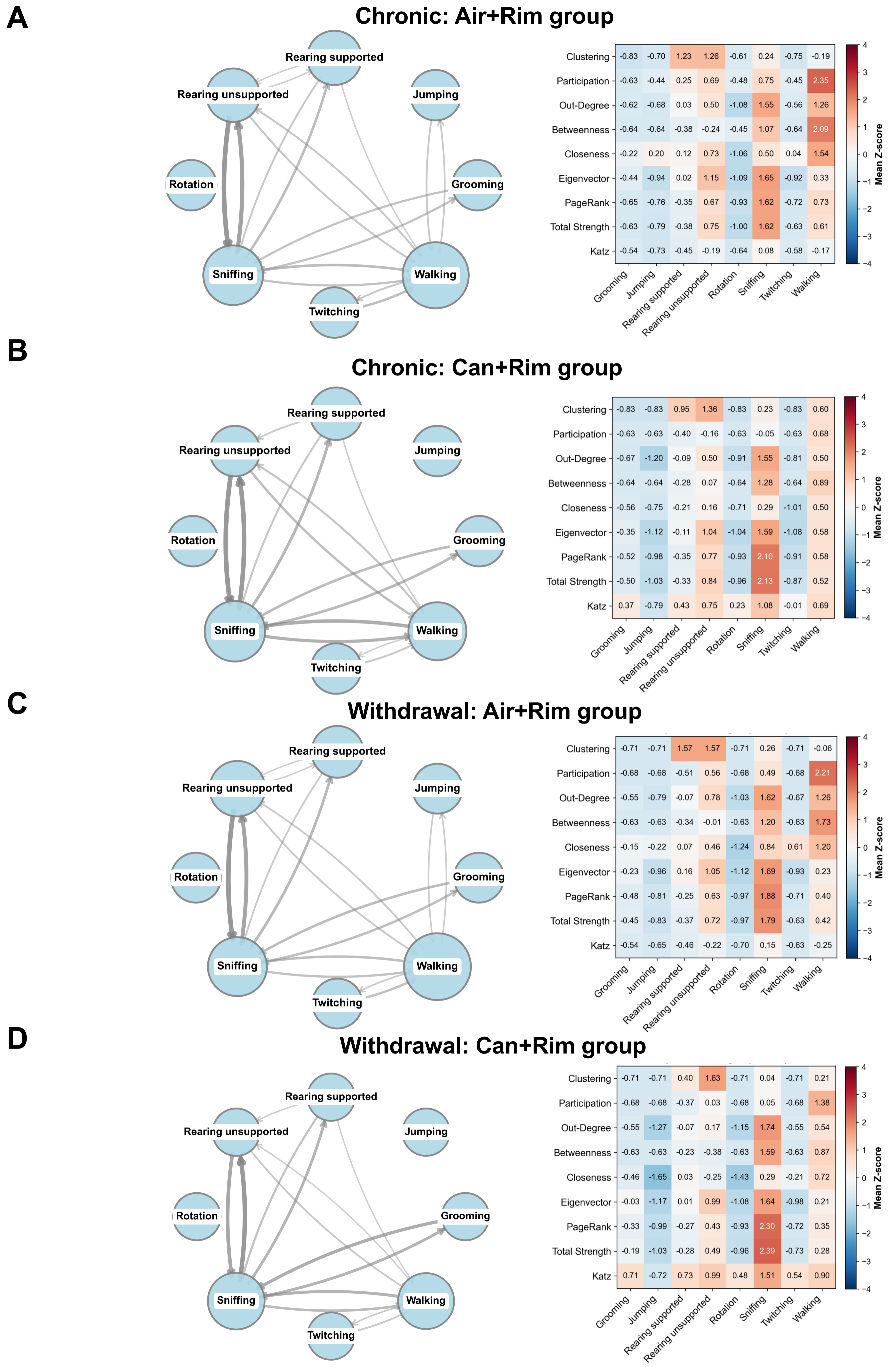

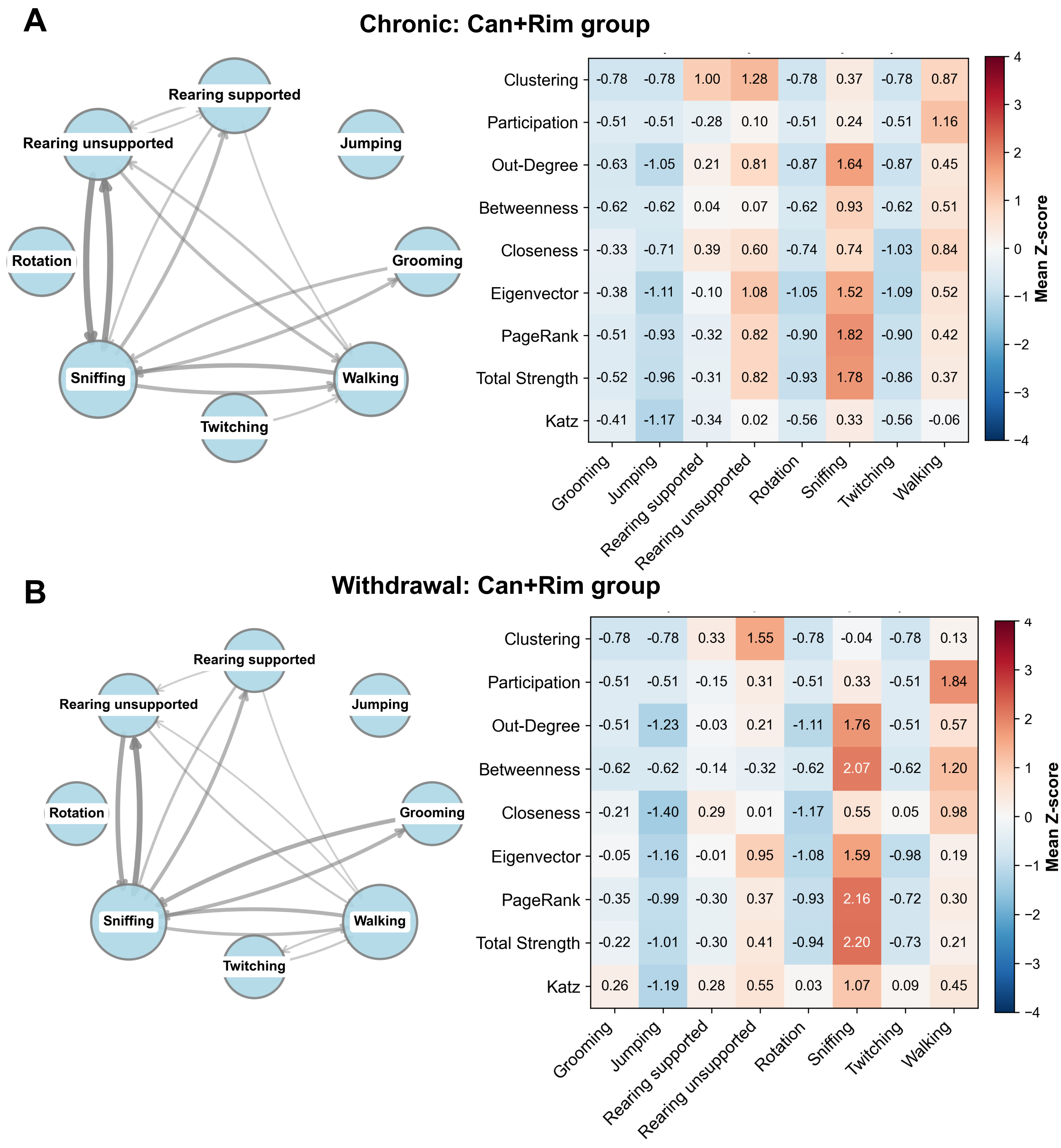

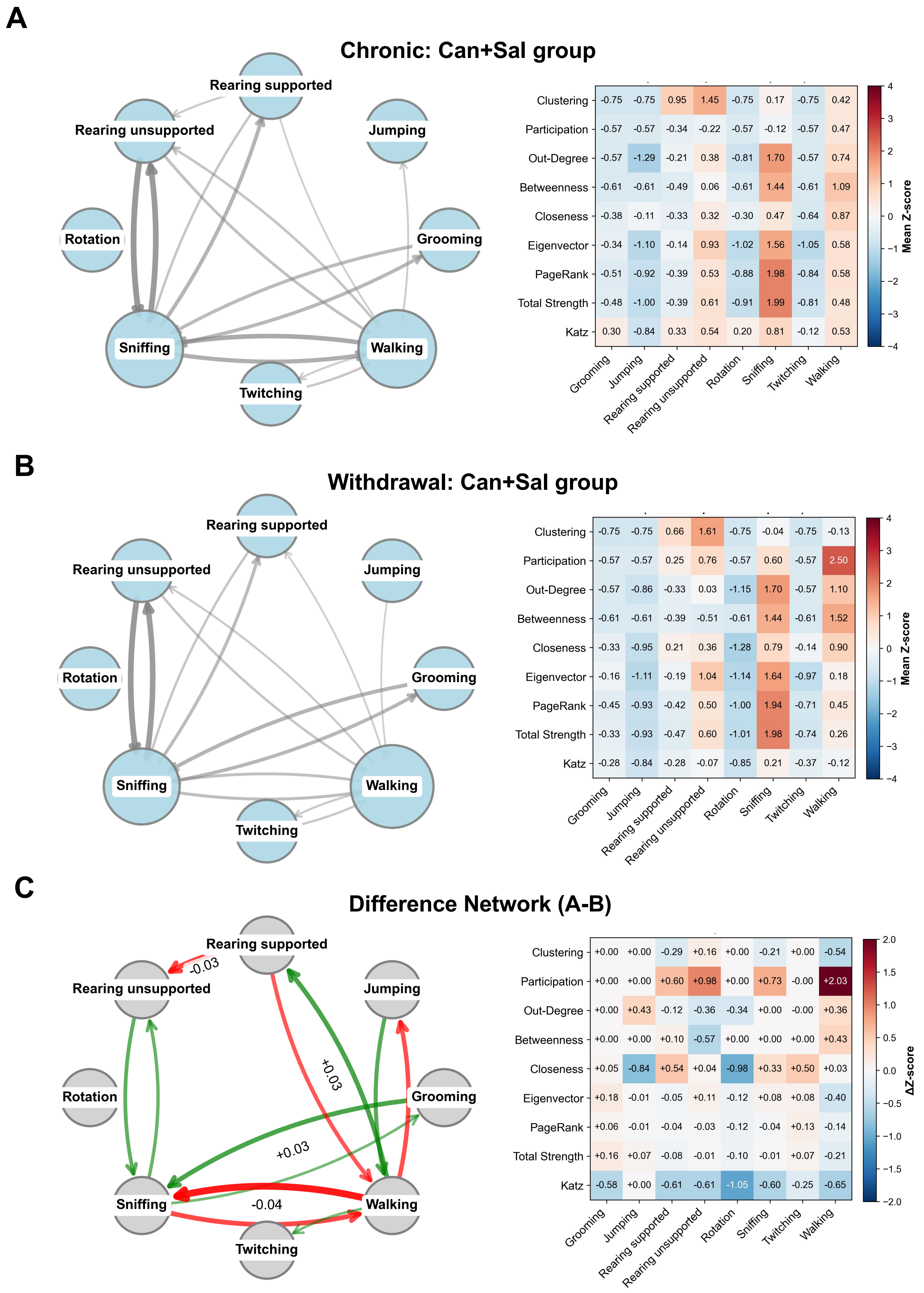
